# The NuRD complex shapes RNA polymerase III activity at highly expressed tRNA gene clusters, tuning the dynamic range of cellular tRNA pools

**DOI:** 10.64898/2025.12.12.694012

**Authors:** Ruiying Cheng, K C Rajendra, Simon Lizarazo, Junmeng Yuan, Sihang Zhou, Kevin Van Bortle

## Abstract

RNA polymerase III (Pol III) produces noncoding RNAs involved in diverse cellular activities, including translation (tRNA, 5S rRNA, 7SL RNA), RNA processing (U6 snRNA, RPPH1, RMRP), and transcription regulation (7SK snRNA). In this way, Pol III activity must be broadly coupled with cellular demands for protein accumulation and growth, increasing in response to nutrient availability and decreasing during differentiation and exit from proliferation. However, the currently established mechanisms of Pol III regulation remain relatively limited, due in part to the few Pol III-centered protein-protein interaction (PPI) studies performed to date. To address this gap, we first investigated PPIs shared by multiple Pol III subunits to understand the macromolecular interactome of Pol III, with special attention directed at potential regulatory candidates. Our proteomic survey uncovers interactions between Pol III and the NuRD (Nucleosome Remodeling and Deacetylase) complex. Taken further, we show that NuRD localizes to active Pol III-transcribed genes and that its recruitment is Pol III-dependent but nonrandom, with peak occupancy and regulatory hallmarks converging on tRNA gene clusters associated with notably high expression levels. Inhibiting NuRD-associated histone deacetylase function reduces Pol III transcription at these sites, suggesting NuRD restricts Pol III and thereby modulates the global dynamic range of Pol III-derived RNA species. These findings are congruent with the transcriptionally repressive nature of NuRD and bring-to-light a new regulatory mechanism that may couple signaling events and changes in metabolic needs with the dynamic availability of specific tRNA pools.

## Introduction

RNA polymerase III (Pol III)-derived noncoding RNAs are generally described as essential housekeeping molecules integral for basic cellular processes, including translation, RNA stability, and processing^1,2^. However, in humans, many Pol III-derived ncRNAs are highly dynamic and, in specific cases, found to be entirely restricted to one or more tissues^2-8^. The dynamic nature of Pol III-derived ncRNA also extends to recurring patterns observed during tumorigenesis, such that specific cancer-emergent ncRNAs have been shown to enhance cell proliferation, migration, and invasion^9-14^. These signatures point to a sophisticated tapestry of underlying transcriptional and posttranscriptional regulatory mechanisms that must contribute to context- and tissue-specific Pol III patterns.

However, in contrast to the plethora of Pol II-centered transcription factors (TFs) and regulatory paradigms^15^, a surprisingly limited number of Pol III-centered regulatory factors have been established. Instead, Pol III transcription is, at present, understood to be largely dependent on the general TFs responsible for Pol III recruitment and initiation at specific promoter architectures, such as TFIIIA (type 1 promoter), TFIIIC (type 1 and 2), variants of TFIIIB (type 1, 2, 3), and the SNAPc complex (type 3)^16-19^. Multiple growth-related pathways converge on Pol III and these recruitment factors, such as through various posttranslational modifications of TFIIIB (e.g. phosphorylation by CK2^20,21^, putative acetylation by SIRT2^22^) and/or specific Pol III subunits. In addition, Maf1, a highly conserved transcriptional repressor, directly antagonizes Pol III activity in response to serum starvation and other mTOR-related signaling events^23^. Myc, on the other hand, enhances Pol III activity through a similarly broad but, instead, positive role in transcriptional amplification^24^.

Here, we sought to investigate additional mechanisms of Pol III regulation using conventional biochemical approaches. We specifically examined Pol III-centered protein-protein interactions (PPIs) with the expectation that presently unknown regulatory features are likely to include factors that make direct contact with the Pol III complex. Using unbiased proteomic methods to map such PPIs, we further prioritized evidence of macromolecular complex-complex interactions, which serve to increase confidence that individual PPIs are reflective of bona fide interactions *in situ*. This framework establishes a novel connection between Pol III and the NuRD (Nucleosome Remodeling and Deacetylase) complex, a chromatin remodeler primarily linked with transcriptional silencing^25,26^. We find that such interactions are functional in nature, such that NuRD reduces Pol III activity with variable efficacy across the Pol III transcriptome. Among other PPIs and complex-complex interactions, our study also captures widespread interactions between Pol III and various other nuclear factors, hinting towards unexpected forms of crosstalk with nuclear processes. These findings advance a basic understanding of RNA polymerase III regulation and function and carry implications for recurrent molecular signatures observed in cancer.

## Results

### Pol III interacts with nucleoporins, splicing factors, and the NuRD chromatin remodeling and deacetylase complex

The 17-subunit Pol III complex is the largest among the eukaryotic transcription enzymes and includes specific subcomplexes involved in its recruitment, transcription initiation and termination, and recycling^27^. To identify Pol III complex interactions, we chose to explore individual protein-protein interactions (PPIs) captured using co-immunoprecipitation followed by liquid chromatography-tandem mass spectrometry (co-IP LC-MS/MS) for subunits of the RPC3-RPC6-RPC7 heterotrimer, a Pol III sub-complex that directly interacts with TFIIIB and thereby facilitates Pol III recruitment to its target gene promoters (Figure 1a)^28,29^. We find that flag-tagged variants of RPC3 (POLR3C), RPC7α (POLR3G), and RPC7β (POLR3GL) each co-immunoprecipitated with RPC1 (POLR3A), the large subunit involving the core site of Pol III transcription (Figure 1b). We additionally demonstrate through LC-MS that flag-RPC3, flag-RPC7α, and flag-RPC7β pull-down experiments are individually enriched for nearly all Pol III subunits, suggesting ectopic Pol III subunits successfully incorporate into the Pol III enzyme (Figure 1c). This is further reflected through gene ontology (GO) enrichment analysis of pull-down-enriched proteins, with the RNA polymerase III complex (GO: 0005666) ranked among the highest enriched GO terms (Figure 1d, Supplementary Figure 1).

**Figure 1.**
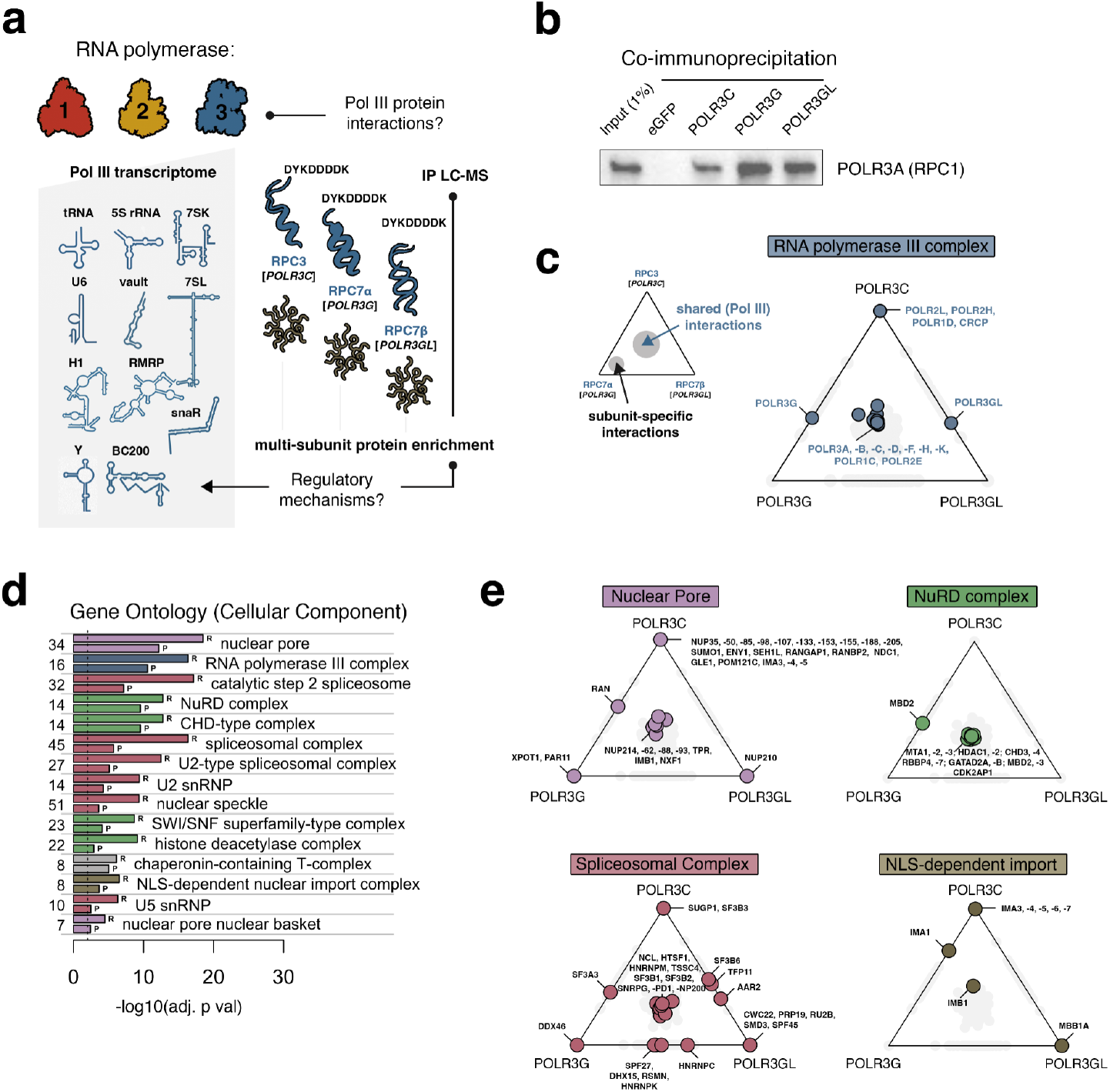
Discovery of protein interactions shared by multiple Pol III subunits identifies putative macromolecular complex interactions. **(a)** Illustrative overview of the unbiased proteomic approach to characterize candidate Pol III protein-protein interactions in the present study. **(b)** co-immunoprecipitation and immunoblot analysis following pull-down of ectopically expressed flag-tagged RPC3 (POLR3C), RPC7*α* (POLR3G), or RPC7*β* (POLR3GL), with large Pol III subunit RPC1 (POLR3A). **(c)** Visualization of subunit-specific and shared protein-protein interactions; schematic illustrates the concept of feature dominance, such that PPIs enriched across all three subunits are centrally positioned. Pol III-complex proteins are enriched in all proteomic experiments, in nearly all cases shared across all three subunits. **(d)** Gene Ontology (cellular component) enrichment analysis of Pol III interactome. GO analysis background: active genes defined by HEK293T RNA sequencing (R) or proteomics (P). **(e)** Analogous visualization of proteins related to specific GO terms enriched in RNA Pol III pull-down experiments, including Nuclear Pore, NuRD complex, Spliceosomal complex, and NLS (Nuclear Localization Signal)-dependent import. n = 3 biological replicates.

In addition to ‘RNA polymerase III complex’, we show that LC-MS experiments are enriched for proteins related to the nuclear pore (GO: 0005643), the U2 spliceosomal complex (GO:0005684), the NuRD (Nucleosome remodeling and deacetylase) complex (GO:0016581), and proteins involved in NLS (nuclear localization signal)-dependent nuclear import (GO:0042564) (Figure 1d). To further visualize the representation of proteins ascribed to enriched GO terms, we transformed the individual enrichment scores for each pull-down experiment into a three-way feature dominance plot. This framework applies a center-of-mass calculation, such that subunit-specific enrichments are drawn to a variable-defined vertex, whereas subunit-shared enrichments are positioned more centrally (Figure 1c)^30^. We note that for Pol III subunit enrichment, RPC7α and RPC7β, which are paralogous and mutually exclusive subunits^31,32^, both RPC7 variants are enriched following RPC3 pull-down, but not for experiments with the mutually exclusive subunit, exemplifying scenarios in which a given protein is enriched in only 2 out of 3 interaction maps (Figure 1c).

Using this approach, we find several nuclear pore (NUP) proteins enriched for all three subunits, including NUP62, NUP88, NUP93, NUP214, and TPR - a nuclear basket protein (Figure 1e). These findings are congruent with previous work identifying Pol III-nuclear basket interactions in C. elegans^33^. RPC3 is specifically enriched for numerous other pore proteins, suggesting extensive interactions with the nuclear pore. This observation may also relate to the RPC3-specific interactions captured with NLS-dependent nuclear import machinery, consistent with the presence of NLS signals only in RPC3 (Figure 1e)^34^. In contrast, the majority of spliceosomal and NuRD-related PPIs are enriched in pull-down experiments for all three subunits, suggesting Pol III establishes macromolecular interactions with specific mRNA splicing and chromatin remodeling machineries (Figure 1e). We hereafter sought to further investigate the functional implications of these Pol III interactions, focusing on the NuRD complex as a priority regulatory candidate.

### The NuRD complex localizes to Pol III-transcribed genes

Given the intuitive implications of NuRD, a multi-faceted complex with roles in gene regulation (Figure 2a)^35,36^, at Pol III-transcribed genes, we directed our primary attention at further establishing NuRD-Pol III interactions and its potential recruitment to canonical Pol III loci. First, we note that independent co-IP experiments for POLR3A - the core Pol III subunit - confirms protein-protein interactions among all NuRD subunits tested (Figure 2b). Likewise, reciprocal co-IP experiments for HDAC1 confirms interactions with several Pol III subunits, including RPC1, RPC2, RPC3, RPC4, and RPC5 (Figure 2c). These experiments, which otherwise fail to capture analogous PPIs with Pol II subunits RPB1, RPB2, RPB3, or Pol I subunit RPA1 (Figure 2d), support a role for NuRD as a macromolecular partner and putative Pol III regulator.

**Figure 2.**
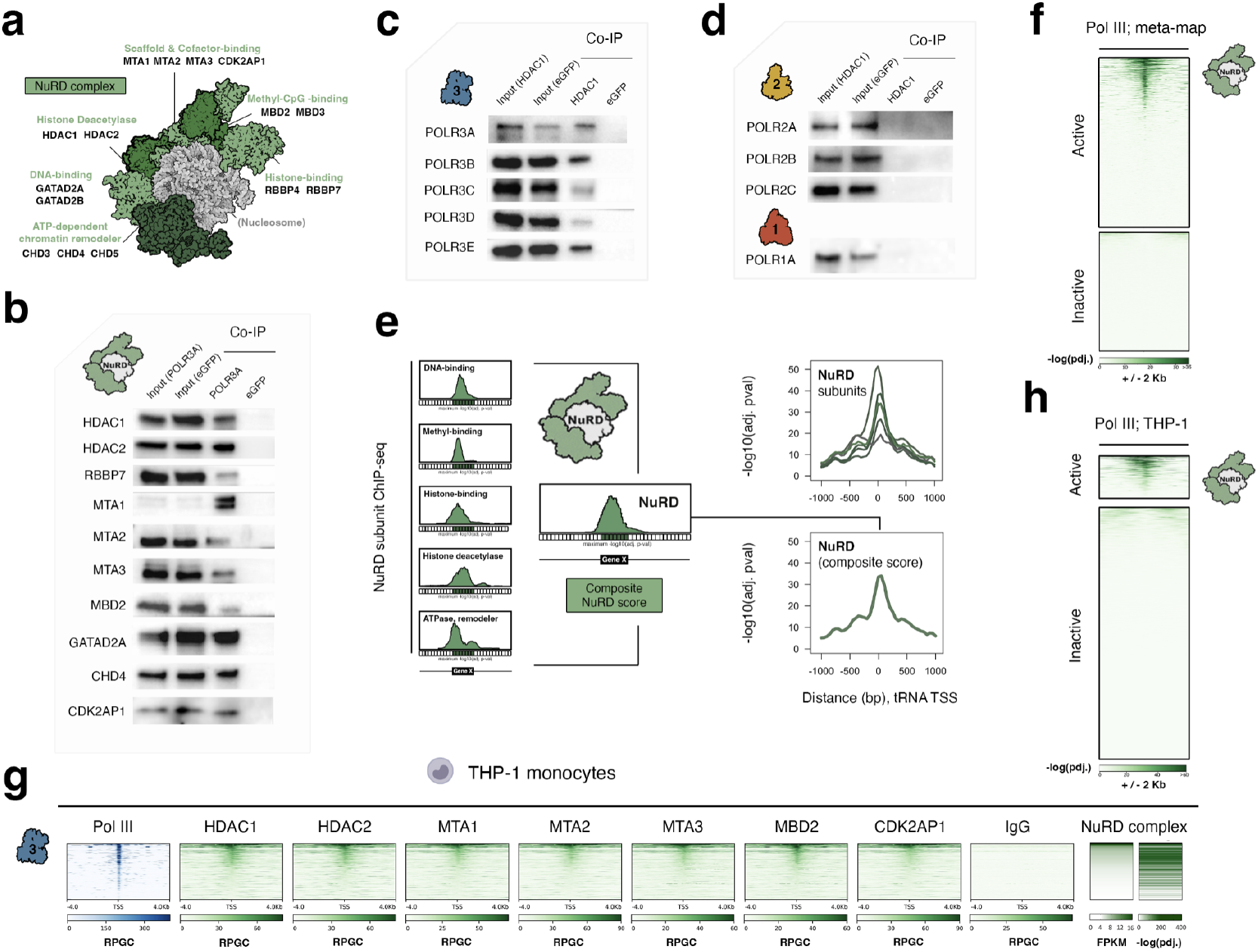
The NuRD chromatin remodeling complex co-localizes with RNA polymerase III at Pol III-transcribed genes. **(a)** Illustrative over-view of the multifaceted NuRD complex, comprised of several interchangeable subunits, structure based on PDB: 7AOA. **(b)** Co-immunoprecipitation and immunoblot analysis for a multitude of NuRD complex subunits following pull-down of ectopically expressed flag-tagged RPC1 (POLR3A), the large Pol III subunit. **(c)** Reciprocal co-IP and immunoblot analysis of Pol III subunits POLR3A, POLR3B, POLR3C, POLR3D, and POLR3E following pull-down of NuRD subunit HDAC1. **(d)** Analogous co-IP analyses for Pol II subunits POLR2A, POLR2B, and POLR2C, and Pol I subunit POLR1A, following pull-down of HDAC1. **(e)** A composite NuRD scoring framework that integrates all currently available ChIP-seq data for NuRD subunits and resulting NuRD occupancy scores at Pol III-transcribed tRNA genes. **(f)** Heatmap analysis of composite NuRD scores across all classes of Pol III-transcribed genes, classified as either “active” or “inactive” across human tissues. **(g)** Heatmap of ChIP-seq signals for the Pol III complex (blue; median read per genomic content (RPGC) of POLR3A, -3B, -3C, -3D, -3E, -3G) and individual NuRD subunits profiled in THP-1 cells across annotated Pol III-transcribed genes. far right: median NuRD occupancy (FPKM) and significance score (green; -log10 adj. p val), sorted by NuRD occupancy. n = 2 biological replicates. **(h)** Heatmap of composite NuRD scores at Pol III-transcribed genes defined as active or inactive specifically in THP-1.

Next, to examine the genomic landscape of NuRD and its positional relationship to Pol III-transcribed genes, we sought to integrate all currently available ChIP-seq data corresponding to individual NuRD subunits. In brief, given the heterogeneity of NuRD subunit composition and the possibility of both complex- and protein-specific binding patterns, we stratified ChIP-seq enrichments by each structural and functional subcategory of NuRD, and thereafter computed a composite NuRD binding score that reflects an overall assessment of NuRD occupancy (Figure 2e). Using this framework, we uncover significant NuRD enrichment at Pol III-transcribed genes, with peak signal intensities located downstream of annotated Pol III start sites (Figure 2e-f). Building further on these insights, we additionally performed ChIP-seq experiments for a multitude of NuRD subunits in THP-1 monocytes, a specific cell line with extensive profiles for Pol III patterns^37-40^. These context-specific experiments uncover widespread overlap of all examined NuRD subunits at Pol III-transcribed genes and, more specifically, Pol III-bound genes, further establishing a direct relationship between NuRD and Pol III at relevant genomic loci (Figure 2g-h).

### NuRD-bound Pol III loci are linked with high expression, positioned nucleosomes, and DNA contact insulation

The co-localization patterns of NuRD at Pol III-transcribed genes indicates that NuRD functions in the context of transcriptionally active rather than inactive loci. However, NuRD is not universally present at all active Pol III elements (defined by Pol III occupancy and production of nascent RNA^38^), suggesting NuRD localization is non-random with respect to the Pol III transcriptome (Figure 2f-h). Perhaps paradoxically, we find that genes with evidence of NuRD-binding are charact-erized by higher accessibility (ATAC-seq), nascent RNA levels (PRO-seq), as well as by higher histone H3K27 acetyl-ation and Pol III occupancy (ChIP-seq; Figure 3a). These data suggest that NuRD recruitment is ineffective at fully repressing Pol III activity at the 215 NuRD-bound elements, despite similar ChIP-seq signatures observed for Pol III and NuRD at these loci (Figure 3b). Even so, it is important to note that numerous highly-expressed and essential Pol III sites are characterized by relatively little NuRD signal, including an array of tRNA gene clusters on chromosome 6 that are known to contribute to the production of critical tRNA pools^41^, in stark contrast to other tRNA gene clusters characterized by more typical NuRD occupancies (e.g. tRNA cluster on chromosome 1; Figure 3c).

**Figure 3.**
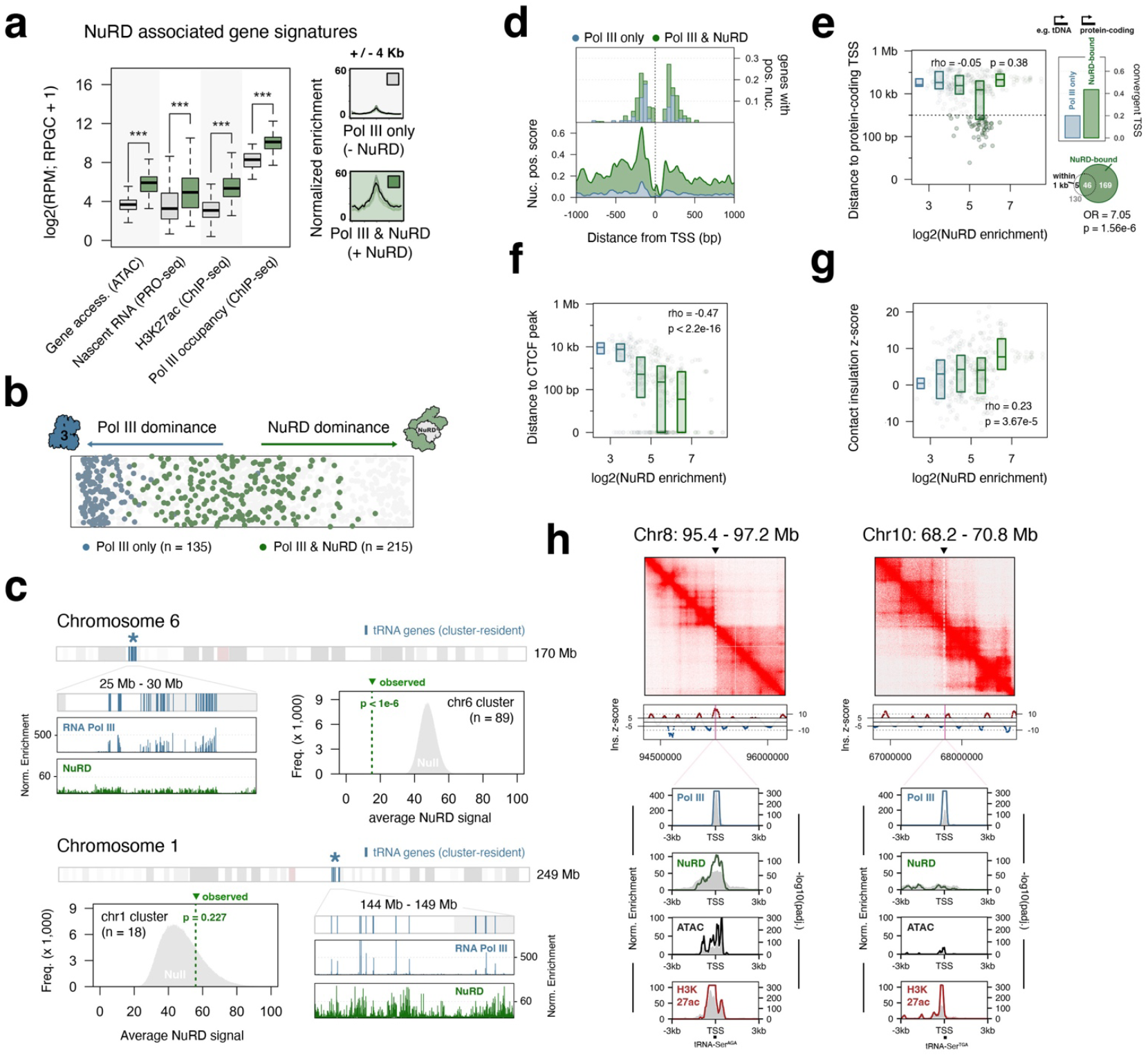
NuRD-occupied Pol III-transcribed genes are active loci associated with hallmarks of nucleosome positioning and chromatin insulation. **(a)** ATAC-seq, PRO-seq, and H3K27 acetylation and Pol III ChIP-seq levels at Pol III transcribed genes that are NuRD-bound (green) or NuRD-free (gray). **(b)** Gene-level enrichment “dominance” analysis comparing Pol III and NuRD-binding signatures. (blue: Pol III only; green: Pol III & NuRD; grey: inactive Pol III genes). **(c)** Example tRNA gene clusters with and without significant NuRD occupancy. Signals denote median normalized occupancy; histograms depict an empirical cluster-level null expectation. **(d)** Nucleosome positioning at active Pol III transcribed genes. top: percentage of genes with positioned nucleosome; bin size 50 bp. bottom: nucleosome position scores. **(e)** Distributions of genomic distance between Pol III genes and protein coding genes (TSS) at increasing levels of NuRD enrichment. Venn diagram depicts the overlap between NuRD-bound genes and those within 1 kb of a protein coding TSS. Right inset shows the percentage of NuRD-bound and NuRD-free Pol III genes with a convergent protein-coding TSS. **(f)** Analogous distribution plots of genomic distance to CTCF-binding sites, at increasing levels of NuRD enrichment. **(g)** Distribution of contact insulation scores, measured by Hi-C, at increasing levels of NuRD enrichment. **(h)** examples of NuRD-bound (left, tRNA-Ser-AGA) and NuRD-free tRNA genes (right, tRNA-Ser-TGA). Top: Hi-C contact frequency heatmap; dotted line indicates tRNA gene coordinates. middle: contact insulation z-score. bottom: Pol III and NuRD ChIP-seq, ATAC-seq and H3K27ac ChIP-seq normalized enrichment (grey area plots) and significance score (colored line plots). Spearman’s rank correlation (ρ, two-tailed P-value).

Given that NuRD-bound Pol III-transcribed genes are transcriptionally active, we surveyed additional chromatin-relevant features at these elements to better understand the possible role or influences of NuRD occupancy. First, we considered nucleosome positioning, as it relates to NuRD-mediated chromatin remodeling^42^. Comparing NuRD-bound to NuRD-free elements reveals significantly higher nucleosome position scores at sites with NuRD, confirming differences in chromatin architecture at these genes (Figure 3d). Although NuRD enrichment scores do not empirically correlate with the genomic distance to protein-coding genes, we find that NuRD is significantly overrepresented at Pol III-transcribed genes that exist within 1 kb of a protein-coding transcription start site (TSS; Figure 3e). Moreover, these sites are nearly twice as likely to be linearly organized in a convergent orientation when NuRD is present, suggesting NuRD recruitment may help insulate Pol III and Pol II activities at gene-proximal loci (Figure 3e). We note that previous work identified a genomic-proximity relationship between NuRD and the architectural protein CTCF, suggesting interplay between NuRD, CTCF, and cohesin distributions and chromatin insulation^43^.

To more directly consider the possibility of chromatin insulation at NuRD-bound Pol III-transcribed genes, we explored the nature of both CTCF-binding and long-range DNA contact insulation in the same cellular context. In brief, we find that NuRD enrichment negatively correlates with the genomic distance to CTCF-binding sites, such that genes with strong NuRD enrichment are likely to exist within 1 kb of a CTCF-bound site (Figure 3f). These data are further congruent with our observation that NuRD enrichment is positively associated with DNA contact insulation, a result that is not otherwise observed for Pol III occupancy alone (Figure 3g, Supplementary Figure 2). Examination of individual loci provides support for this large-scale trend, with examples of NuRD-occupied Pol III-genes existing at the boundaries of topologically-associated domains (TADs), in contrast to genes dominated by Pol III and absent NuRD binding (Figure 3h). Nevertheless, the convergence of NuRD with CTCF, a well-established insulator protein, confounds whether NuRD actively contributes to chromatin insulation at these sites^44-47^.

### NuRD-Pol III co-localization peaks at specific tRNA gene clusters within a variable number tandem repeat (VNTR)

The nonrandom organization of NuRD at specific Pol III-transcribed gene clusters (examples in Figure 3c) led us to further examine NuRD occupancy with respect to all linear clusters of tRNA genes and other elements. In brief, the human Pol III transcriptome is to a large degree organized into physically proximal clusters of Pol III-target genes^39,48^. Considering the maximum NuRD ChIP-seq signal intensities, we find that NuRD patterns peak at specific tRNA gene clusters encoded within a variable number tandem repeat (VNTR) on chromosome 1 (Figure 4a)^49^. To more broadly consider and compare NuRD-binding at all Pol III-target gene clusters, we applied a global clustering algorithm that groups all individual genes encoded within an aggregative, 20 kb maximum gene-to-gene distance (Figure 4b). Using this clustering approach, we find that the VNTR-encoded tRNA gene clusters are significantly enriched for NuRD, far more than for all other Pol III-target gene clusters (Figure 4b, Supplementary Figure 3), suggesting NuRD localization may play a distinctly important role at VNTR-associated tRNA genes.

**Figure 4.**
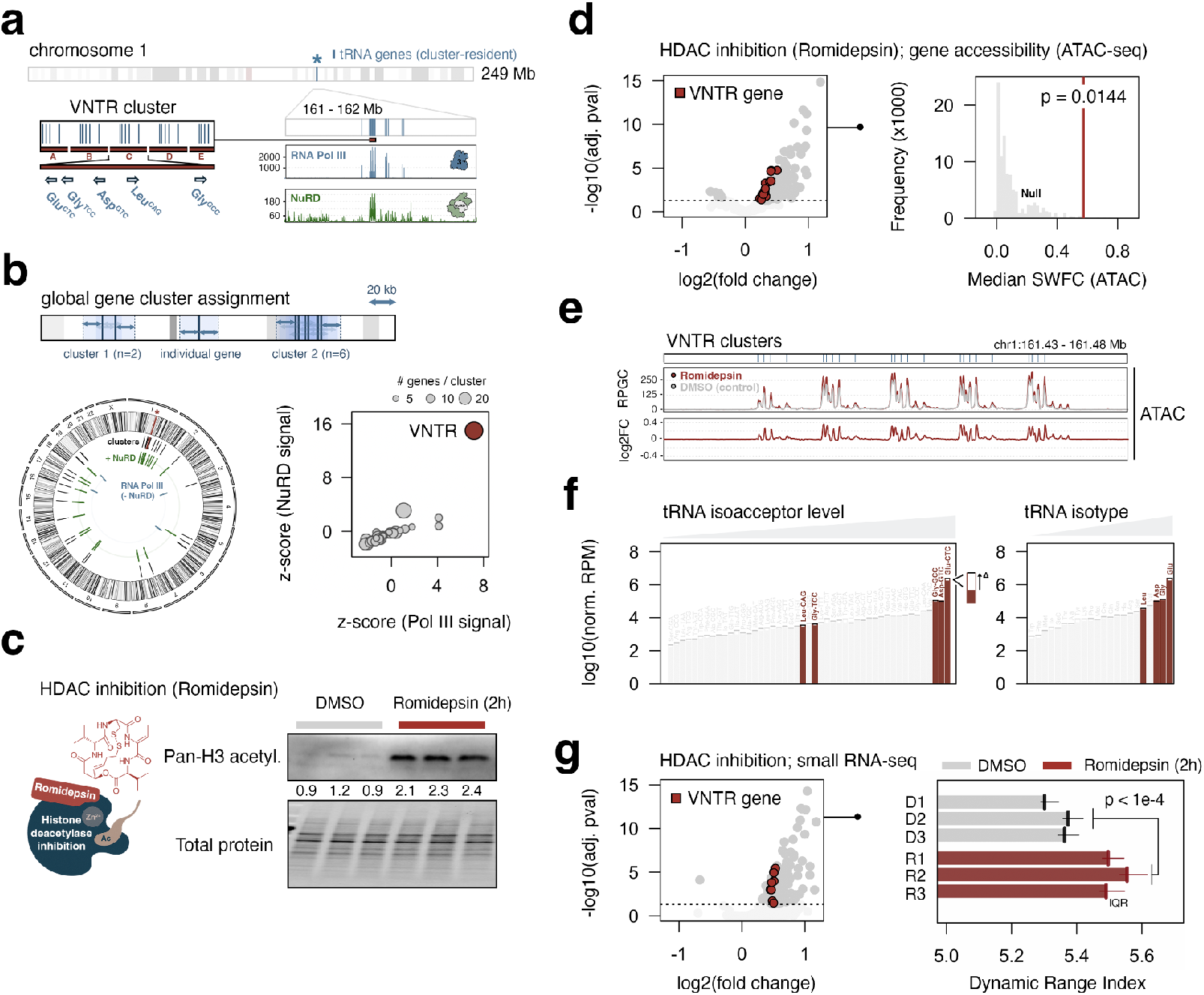
NuRD histone deacetylase function restricts Pol III transcription at highly expressed tRNA gene clusters, thereby shaping the dynamic range of the Pol III transcriptome. **(a)** VNTR-encoded tRNA gene organization and ChIP-seq enrichment for Pol III (blue) and NuRD (green). Blue dashes denote active, cluster-proximal tRNA genes. **(b)** Illustration of agglomerative gene clustering method (top) and resulting Pol III target gene clusters (bottom left); Pol III and NuRD enrichment z-scores (bottom right) quantify the cluster-level significance of Pol III and NuRD enrichment, relative to an empirical null expectation. **(c)** Chemical structure and mechanism-of-action of Romidepsin, a class I HDAC inhibitor (left). Immunoblot analysis of Histone H3 acetylation and total protein levels in control (DMSO) and Romidepsin-treated THP-1 cells (p=0.0007; Student’s t-test). **(d)** differential gene accessibility profile across active Pol III-transcribed genes; genes with FDR <.05 highlighted in dark grey (VNTR genes highlighted in red). Observed significance-weighted fold change (SWFC) for ATAC-seq at VNTR-encoded tRNA genes, compared to an empirical null distribution (p=0.0144; permutation test). **(e)** ATAC signal and log2(fold change) at VNTR cluster. **(f)** Comparison of tRNA abundance in control (DMSO) and romidepsin-treated cells, at both isoacceptor-level and isotype-levels. Black bars indicate directional change; VNTR tRNA highlighted in red. **(g)** differential small RNA-seq analysis for Pol III-transcribed genes following romidepsin treatment (left) and global dynamic range index comparison between control (DMSO; gray) and romidepsin-treated cells (red; p-value < 1e-4; permutation test).

### NuRD function antagonizes highly expressed Pol III genes and limits the dynamic range of cellular tRNA pools

The NuRD complex integrates ATP-dependent chromatin remodeling with targeted histone deacetylation to exert its transcriptionally repressive functions^50,51^. Given that the redundancy of both chromatin remodeling (CHD3, CHD4, CHD5) and histone deacetylase subunits (HDAC1 and HDAC2) presents a challenge with respect to effectively silencing NuRD activity, we relied on pharmacological inhibition of NuRD deacetylase function via Romidepsin (FK 228^52^) - a potent and selective inhibitor of HDAC1 and HDAC2 - to broadly restrict NuRD-mediated deacetylation. We specifically investigated the effect of Romidepsin (50 nM, IC50 HDCA1=36 nM, IC50 HDAC2= 47 nM) on Pol III activity following a 2-hour exposure, which is sufficient at significantly increasing the total histone H3 acetylation levels in THP-1 monocytes (Figure 4c). We find that HDAC1/2 inhibition significantly increases the chromatin accessibility at Pol III-transcribed genes, including at VNTR-associated tRNA genes enriched for NuRD occupancy (Figure 4d-e). Taken as a group, the increase in accessibility at VNTR-associated genes is statistically higher than expected across all Pol III-transcribed genes, consistent with the expectation that NuRD plays a notable role at this locus (Figure 4d, p = 0.0144, permutation test).

The dynamic accessibility of VNTR-associated tRNA genes following HDAC inhibition is consistent with evidence that NuRD can restrict chromatin accessibility, in contrast to other chromatin remodeling complexes that generally promote DNA accessibility^53^. Given that changes in accessibility ostensibly modulate transcription of NuRD-associated genes, we further integrated ATAC-seq experiments following HDAC inhibition with small RNA-seq analysis of Pol III-derived ncRNA species. Expression of VNTR-associated genes, which include those encoding the three most abundant tRNA isoacceptors and four out of the five most abundant tRNA isotypes, increases significantly in response to the 2-hour Romidepsin exposure (Figure 4f-g), consistent with observed changes in gene accessibility.

Beyond the locus-dominant enrichment of NuRD and evidence that NuRD antagonizes select Pol III-transcribed genes, the VNTR-associated tRNAs are notable for defining the high-end of cellular tRNA pools. In this way, these elements effectively set the dynamic range of the Pol III transcriptome, which may stretch to variably low, moderate, or high levels contingent on both transcriptional and post-transcriptional regulatory mechanisms^54^. The dynamic abundance of such tRNA pools are functionally relevant, as highly expressed tRNAs have evolutionarily adapted to high frequency “optimal” codons, which are themselves enriched in mRNAs encoding growth-related proteins and influence translation efficiency^55^. To better quantify the Pol III-derived small RNA dynamic range (independent of simply measuring the most abundant RNA species), we subsampled the distribution of Pol III products and estimated the median level of the 5 most abundant RNA species. Applying this framework to THP-1 monocytes in the presence or absence of romidepsin, we find that inhibiting NuRD-associated deacetylation indeed significantly increases the dynamic range of cellular tRNA pools (Figure 4g, Supplementary Figure 4). These data suggest a model in which NuRD functionally antagonizes the level of Pol III transcription through direct localization and regulation at highly expressed Pol III target genes, consequently modulating the dynamic range of the Pol III transcriptome.

### NuRD co-occupancy at Pol III-transcribed genes is Pol III-dependent, suggesting direct recruitment and regulation

Our identification of the NuRD complex as a bona fide Pol III interactor and discovery that NuRD localizes to transcriptionally active (rather than inactive) loci indicates that Pol III itself may play a direct role in recruiting NuRD to Pol III-transcribed genes (Figures 1e, 2b-c, 3a). To assess whether NuRD recruitment is at all Pol III-dependent, we first examined the effect of ML-60218 (ML-6), a small molecule Pol III-specific transcriptional inhibitor that also causes Pol III displacement^56,57^, on NuRD-binding patterns. Overall, ML-6 inhibition leads to a significant decrease in ChIP-seq signal at Pol III-target genes for several NuRD subunits (Figure 5a). Moreover, ML-6-dependent loss of NuRD enrichment is most pronounced at VNTR-resident tRNA genes, further suggesting that NuRD recruitment and activity at these select, highly expressed tRNA genes is at least partially dependent on Pol III residency (Figure 5a-b, Supplementary Figure 5). In contrast to HDAC1/2 inhibition, which increased the dynamic range of Pol III-derived ncRNAs, ML-6 reduces Pol III activity and thereby restricts the dynamic range of tRNAs, including via significant reductions in nascent transcription of VNTR-encoded tRNA isoacceptors and isotypes (Figure 5c). While the reduction of Pol III activity at these sites is likely caused primarily by direct Pol III inhibition, we note that comparison of Pol III and NuRD signals before and after ML-6 treatment uncovers a strong shift in gene occupancy “dominance”, such that NuRD increases relative to Pol III, particularly at VNTR-encoded elements, following Pol III inhibition (Supplementary Figure 6). These data indicate that NuRD localization is Pol III-dependent, but that it may also exert a stronger repressive regulatory effect in response to outside challenges to Pol III function.

**Figure 5.**
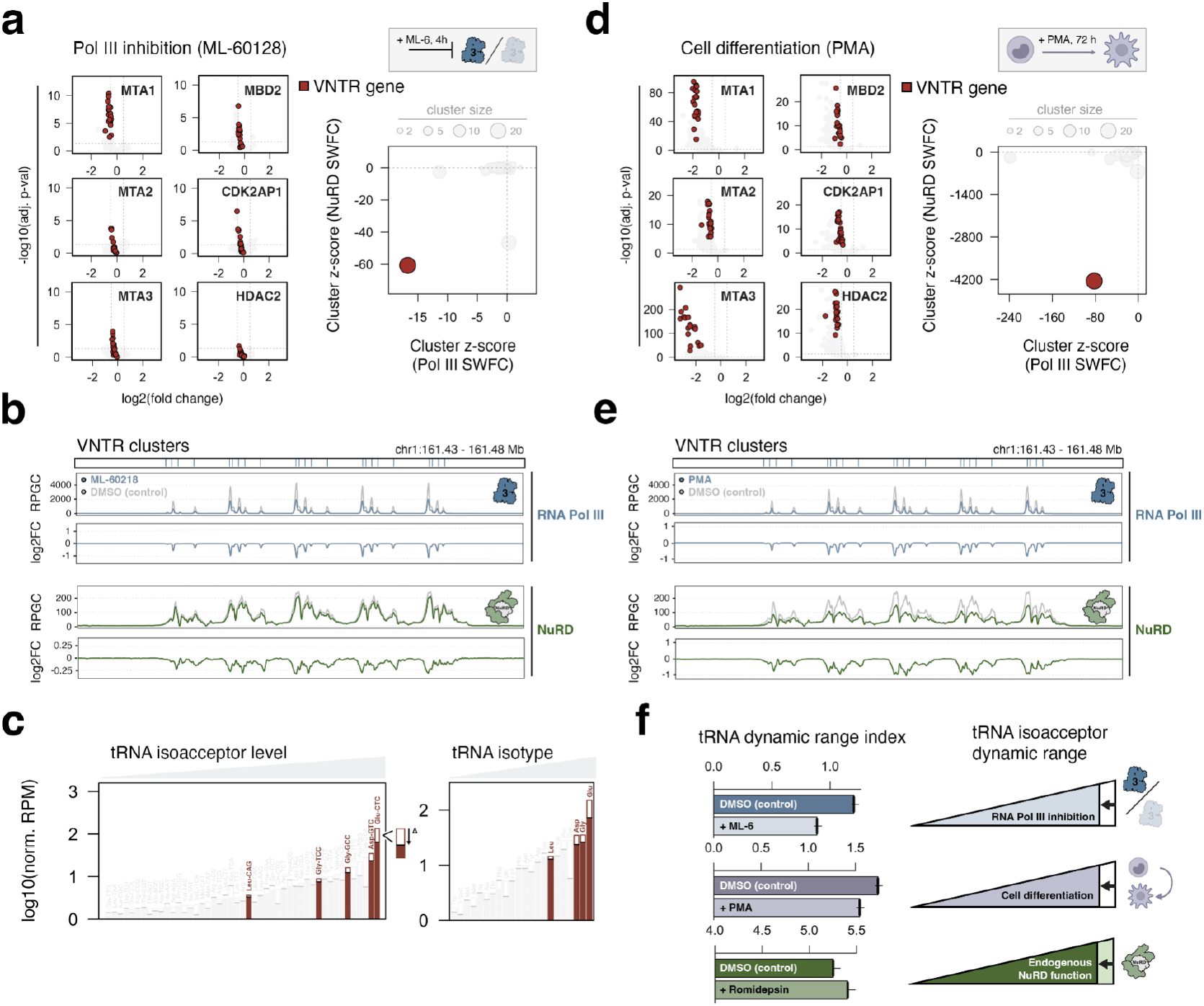
NuRD recruitment to Pol III-transcribed genes is sensitive to Pol III inhibition and to differentiation-associated Pol III down-regulation. **(a)** differential ChIP-seq analysis of NuRD subunit-binding following Pol III-inhibition (ML-60128; left), dynamic Pol III and NuRD z-scores (right) quantify the cluster-level significance of Pol III and NuRD significance-weighted fold change (SWFC) values, relative to an empirical null expectation. **(b)** ChIP-seq signal and log2(fold change) for Pol III and NuRD at VNTR tRNA gene cluster in response to Pol III inhibition. **(c)** Comparison of nascent tRNA abundance in control (DMSO) and ML-60218-treated cells, at both isoacceptor-level and isotype-levels. Black bars indicate directional change; VNTR tRNA highlighted in red. **(d)** differential ChIP-seq analysis of NuRD subunit-binding during THP-1 differentiation (PMA; left), dynamic Pol III and NuRD z-scores (right) quantify the cluster-level significance of Pol III and NuRD significance-weighted fold change (SWFC) values, relative to an empirical null expectation. **(e)** ChIP-seq signal and log2(fold change) for Pol III and NuRD at VNTR tRNA gene cluster following THP-1 monocyte-to-macrophage differentiation. **(f)** Quantitative comparison of tRNA dynamic range indices in response to Pol III inhibition (top), cell differentiation (middle), and HDAC1/2 inhibition (bottom); model positions NuRD as an endogenous regulator of Pol III transcription with important consequences on the dynamic range of Pol III-derived tRNAs and other ncRNAs.

To further examine the effect of Pol III dynamics on NuRD localization, we additionally profiled NuRD subunit binding patterns following PMA-induced THP-1 monocyte differentiation, which leads to widespread downregulation of Pol III occupancy and transcription^38,39^. As observed following Pol III inhibition, we find that NuRD localization is significantly reduced at Pol III-transcribed genes, including at VNTR-encoded elements, concomitant with loss of Pol III during differentiation (Figure 5d-e). While NuRD depletion is again most pronounced at the VNTR locus (Figure 5d, Supplementary Figure 7), we show that differentiation also produces a shift in gene-level dominance, with NuRD increasing in THP-1 macrophages relative to Pol III occupancy (Supplementary Figure 6). These patterns are once again linked with similar reductions in the dynamic range of tRNAs, further mirroring observations following Pol III inhibition (Figure 5f).

## Discussion

By leveraging a Pol III-centered protein-protein interaction discovery framework, our present study demonstrates that Pol III interacts with the NuRD complex, uncovering a previously unrecognized regulatory axis that is altogether supported through both biochemical and large-scale genomic data. For one, ChIP-seq analyses confirm that NuRD is highly enriched at Pol III-transcribed genes, linking our novel protein-protein interactions to well-established sites of Pol III transcription. Taken further, we show that functional inhibition of HDAC1/2 leads to an acute increase in Pol III-derived ncRNA levels, indicating that Pol III-NuRD interactions and co-occupancy at Pol III-transcribed genes involves negative gene-level regulation, consistent with the repressive nature of NuRD function. Though NuRD-binding appears sensitive to both Pol III inhibition- and differentiation-associated gene displacements, we show that NuRD localization is not equally distributed across Pol III-target genes, suggesting Pol III and additional factors may collectively contribute to NuRD recruitment to select genes.

The subset of Pol III-transcribed genes characterized by maximum NuRD enrichment and sensitivity to HDAC1/2 inhibition correspond to multi-copy tRNA genes encoded within a VNTR on chromosome 1. This finding is notable, first of all, for the high level of Pol III transcription at these genes, suggesting proximal density and local enrichment of Pol III may supercharge the recruitment of NuRD to these elements. The suborganization of NuRD to this site also holds implications for the dynamic range of cellular tRNA pools, such that transcriptional antagonism by NuRD may limit the “stretch” from low-to-high abundance tRNA species. Given that high abundance tRNAs correspond to evolutionarily optimal codons enriched in growth-related mRNAs, NuRD-mediated antagonism may conceivably buffer Pol III activity from overactivity, ultimately opposing growth-favoring codon-specific translation efficiencies.

Together, these findings introduce NuRD as a novel component of the Pol III regulatory sphere, in addition to the well-established core transcription factors (e.g. TFIIIA, TFIIIB, TFIIIC, SNAPc) and signal-response proteins (e.g. Maf1, MYC). The deacetylation-dependent role of NuRD is unique among Pol III regulatory factors, however, instead positioning NuRD as a chromatin-associated player that coexists with high local transcription and tempers rather than silences Pol III. The Pol III-dependence of NuRD occupancy interestingly suggests that Pol III attracts yet is restrained by this chromatin-based mechanism.

While our study of Pol III interactions successfully identified NuRD as a priority regulatory candidate, we note that our proteomic survey also uncovered many previously unknown Pol III-centered PPIs, hinting at an extensive network of interactions between Pol III and nuclear pore subcomponents, proteins involved in mRNA splicing, and many additional factors. These data point to currently unidentified forms of crosstalk involving Pol III and other distinct nuclear processes, highlighting the need for deeper exploration of Pol III within the broader lens of genome biology.

## Limitations of this study

Although co-IP mass spec analyses were useful at identifying novel Pol III-centered PPIs, this approach is inherently biased and may fail to identify low-affinity interactions or those that occur with limited frequency. Our framework also prioritized interactions shared across multiple Pol III subunits (POLR3C, POLR3G, POLR3GL and, in the case of NuRD, confirmed by POLR3A). While this decision was made in the interest of boosting confidence of candidate Pol III complex-complex interactions, it is likely that some regulatory factors function through subunit-specific rather than multi-subunit interactions. Additional regulatory mechanisms are likely to involve context-dependent and/or transient interactions as well as indirect, chromatin-mediated associations not captured by our approach. Future use of proximity-labeling or crosslinking-based methods may provide insight on these additional players.

The multi-faceted nature of the NuRD complex and the redundancy within each modular function (chromatin remodeling by CHD3, CHD4, or CHD5; deacetylation by HDAC1 or HDAC2, DNA-binding by MBD2 or MBD3, etc.) presents a unique challenge, both in terms of disentangling the putative role and effect of each independent function and for addressing compensatory mechanisms. Moreover, both biochemical and genomic findings indicate that Pol III interacts and co-localizes with all possible “identities” of the NuRD complex, precluding a targeted experimental analysis. Thus, our decision to focus on deacetylation by pharmacological inhibition of HDAC1 and HDAC2 was predicated on the efficacy of Romidepsin to rapidly inhibit both enzymes. Understanding the contribution of nucleosome remodeling and other features of NuRD activity therefore remain open questions to address.

Finally, while our results indicate that NuRD localization may be Pol III-dependent, the molecular basis for this putative dependency remains unclear. For example, NuRD recruitment may involve physical contact between specific Pol III and NuRD subunits, interactions with Pol III-related ancillary factors, via chromatin modifications and/or nascent RNA molecules produced by Pol III, or, perhaps more likely, through a combination of such factors. These potential interactions highlight the overall complexity and challenge for future studies aimed at further deconstructing the Pol III-NuRD regulatory axis.

## Materials and Methods

### Data acquisition

Raw ChIP-seq data underlying the context-agnostic NuRD metamap were retrieved from the NCBI GEO database (see Supplementary Table 1 for a corresponding accession list). Signal track files corresponding to processed composite NuRD metamap ChIP-seq and uniform bin-scores were generated using deeptools^58^; resulting data are available for download as individual .bigwig files via Zenodo https://zenodo.org/records/13274508. THP-1 nascent small RNA sequencing (PRO-Seq), H3K27ac and CTCF ChIP peaks were obtained from GSE96800 and THP-1 monocyte Pol III ChIP seq are obtained from GSE163422. THP-1 differentiation related small RNA-seq data were obtained from GSE171884. THP-1 In situ Hi-C data were retrieved from SRA bioproject number PRJNA385337

### Cell lines and culture conditions

THP-1 monocytes were obtained from ATCC (TIB-202 Batch# 62454382) and propagated in T-75 flasks between 0.2 and 1 × 10e6 cells/mL in an RPMI-1640 (Catalog# 11875093, Gibco, Billings, MT, USA) growth medium. THP-1 experiments were conducted on cells between passage 10 and 15. THP-1 derived macrophages were collected after 72-h exposure to PMA by aspirating media and any non-adherent cells, and incubating adherent cells with TrypLE (ThermoFisher) for 15 min. THP-1 cells were treated with RNA polymerase III inhibitor ML-60218 (Catalog# 557403-10MG, SigmaAldrich) at a final concentration of 25 µM; Romidepsin(FK228;Catalog#HY-15149, MedChemExpress) at a final concentration at 50 nM and cells collected at 2 h post-exposure followed by cell wash in PBS buffer.

HEK293T cells were obtained from ATCC (CRL-3216, Batch#70049877) and grown in 10 cm BioLite™ Cell Culture Treated Dishes (Catalog# 12-556-002 Thermo Scientific) in Dulbecco’s Modified Eagle Medium, high glucose (Catalog# 11965092, Gibco). HEK293T experiments were conducted on cells between passage 10-25.

### Plasmid and Antibody reagents

Sequence-verified pcDNA3.1 + /C-(K) DYK (Flag) plasmids encoding eGFP, POLR3A (Clone ID: OHu27038D), POLR3C (Clone ID: OHU31224D), POLR3G (Clone ID: OHU054866C), PLOR3GL (Clone ID: OHUU30091D) and HDAC1 (Clone ID: OHu27230) were obtained from GenScript.

Antibody reagents used in this study included anti-MTA3 (Proteintech, #14682-1-AP), MTA2 (Invitrogen, #PA1-41581), MTA1 (Proteintech, #30545-1-AP), MBD2 (Proteintech, #55200-1-AP), HDAC2 (ActiveMotif AbFlex, #91197), HDAC1 (Proteintech, #10197-1-AP), CDK2AP1 (Proteintech, #13060-2-AP), POLR2A (Proteintech, Cat No. 20655-1-AP), POLR2B (Proteintech, Cat No. 20370-1-AP), POLR2C (Proteintech, Cat No. 13428-1-AP), POLR1A (Proteintech, Cat No. 20595-1-AP), POLR3A (Thermo Fisher, Cat No. PA5-58170), POLR3B (Bethyl, Cat No. A301-855A), POLR3C (Bethyl, Cat No. A303-063A), POLR3D (Bethyl, Cat No. A302-295A), POLR3E (Bethyl, Cat No. A303-708A), Histone H3ac (pan-acetyl) (Active Motif, Cat No. 39140).

### Immunoblotting and protein quantification

Cell pellets were washed once with PBS before lysis with RIPA buffer (Catalog# J62524.AD, Thermo Scientific, Waltham, MA, USA) following standard protocols. Total protein concentration was determined using a Pierce BCA protein assay kit (Catalog# 23225, Thermo Scientific, Waltham, MA, USA), equivalent protein fractions, diluted in RIPA buffer are incubated at 95°C with diluted 4× Laemmli Sample Buffer (Catalog#1610741, BIO-RAD Hercules, CA, USA) for 5min. Proteins were separated on 4– 20% Mini-PROTEAN® TGX Stain-Free™ Protein Gels, 15 wells (Catalog# 4568096, BIO-RAD, Hercules, CA, USA) using 10× Tris/Glycine/SDS (Catalog# 1610732, BIO-RAD, Hercules, CA, USA) and transferred onto polyvinylidene difluoride membranes (0.2 um) (Catalog# LC2002, Invitrogen, Waltham, MA, USA) with a Trans-Blot® Turbo™ Transfer System (Catalog# 1704150, BIO-RAD, Hercules, CA, USA). Transfer membranes were blocked with a 5% blotting-grade blocker (Catalog# 1706404, BIO-RAD, Hercules, CA, USA), followed by incubation with the primary antibody at 4 °C overnight. Membranes were washed with TBST and incubated with mouse or rabbit secondary antibodies conjugated with horseradish peroxidase (Catalog# 31430, 31462, Invitrogen, Waltham, MA, USA) for 2 h at room temperature followed by three washes in TBST. Proteins were visualized using either SuperSignal West Pico (Catalog# 34580, Thermo Scientific, Waltham, MA, USA) or SuperSignal West Femto (Catalog# 34096, Thermo Scientific, Waltham, MA, USA) with a ChemiDoc™ Touch Imaging System (Catalog# 1708370, BIO-RAD, Hercules, CA, USA). Protein abundances were calculated by normalizing the total protein intensity per lane.

### CoIP LC-MS/MS

Flag-tagged NuRD subunit HDAC1, Pol III subunits RPC3 RPC7 RPC7l and eGFP control are overexpressed in HEK293T cells using Lipofectamine 3000 Transfection Reagent (Cat #L3000015) following manufacturer’s protocol. Cells are collected 2 days after transfection and coIP performed using Pierce™ Classic Magnetic IP/Co-IP Kit(88804) and Pierce™ Anti-DYKDDDDK Magnetic Agarose(A36797) following manufacturer’s protocol. The samples are subjected to mass-spectrometry by nano-LC-MS/MS (Ultimate 3000 coupled to a Q-Exactive HF, Thermo Scientific) at the University of Illinois Carver proteomics core. The mass spectrometry proteomics data have been deposited to the ProteomeXchange Consortium via the PRIDE73partner repository with the dataset identifier PXD061197 and 10.6019/PXD061197

### Mass spectrometry data analysis

For sample-sample normalization, the intensities of each protein (eGFP(n=3), POLR3C(n=3), POLR3G(n=2), POLR3GL(n=3)) were divided by sample-total intensity. Non-specific binding proteins are filtered out by removing any proteins in any of the three eGFP pulldowns. The average signal intensity is calculated for each binding protein. The enrichment scores are calculated using log2[(ave signal intensity+1*10^-8)/ 1*10^-8]. HEK293T-specific GO analyses were performed considering all proteins enriched within one or more bait Pol III subunits using DAVID Knowledgebase v2024q2, using either the HEK293T transcriptome (GSE268457) with lower threshold of TPM>0, or HEK293T proteome (PXD055286), as the expectation universe for the category *GOTERM_CC_ALL*. Significance testing involved one-sided hypergeometric tests, with resulting p-values adjusted using the Benjamini-Hochberg (FDR) method.

### ATAC-seq and Nucleosome positioning analysis

ATAC seq samples of DMSO and 50nM Romidepsin treated THP-1 cells were prepared from ∼70k cells per sample using ATAC-Seq Kit (Cat #53156 Active Motif), following the manufacturer’s protocol for Cell Sample Preparation. Libraries were quantified by bioanalyzer and sequenced on a NovaSeq X Plus on a 25B lane with 2×150nt reads.

Fastq files were generated and demultiplexed with the bcl2fastq v2.20 Conversion Software (Illumina). ATAC-seq reads were trimmed using Trim_Galore version 0.6.5 (https://github.com/FelixKrueger/TrimGalore); aligned to genome reference GRCh38 using Bowtie2 version2.3.5.1^59^; and sorted & indexed using SAMtools version 1.9^60^. All reads less than 20 bp, unmapped, and/or aligned to the mitochondrial genome were filtered. Read duplicates were filtered using GATK picard version 2.9.4^61^. Individual sequencing replicates were thereafter uniformly scaled to 120 million reads using the samtools view -s function. Read counts were extracted over currently annotated ncRNA genes – retrieved from RNAcentral^62-64^ (gene body +-150bp) using bedtools coverage -counts (BEDTools version 2.26^65^). ATAC-seq peaks were called using MACS2 version 2.2.5^66-69^ and differential analysis performed using DEseq2^70^.

Nucleosome position analyses were performed with NucleoATAC version 0.3.4^71^. Positioned nucleosomes called in nucmap_combined.bed were retained if (a) # of reads in 121 bp window > 2 and (b) occupancy > 0.2. To further increase confidence, we considered only genes with an ATAC-seq peak called in all three replicates. For each gene, the +/-1 nucleosome was called in a 1kb window upstream of the TSS or 1kb downstream of TES. Genes with +/-1 nucleosome dyed consistently within 200 bp in all three replicates are marked as “positioned nucleosome positive”. Bin-wise nucleosome occupancy scores were extracted from *.nucleoatac_signal.smooth.bedgraph; directionalities were considered before extracting bin-mean across active Pol III genes. The ratio of genes with a positioned nucleosome were defined as # NuRD bound(unbound) genes with an ATAC-seq peak and positioned nucleosome / # NuRD bound(unbound) genes with an ATAC-seq peak.

### Global NuRD occupancy scoring (NuRD metamap)

We leveraged ChIP-seq data from GEO and ENCODE datasets for CHD4, GATAD2A, GATAD2B, HDAC1, HDAC2, MBD2, MBD3, MTA1, MTA2, RBBP4 respectively (Supplementary Table 1). We implemented a comprehensive normalization procedure described in K C et. al^30^. Briefly, for each dataset, read counts were extracted across 50 bp genome bins, then normalized to 100 million total read counts, then sums are calculated for each bin as the observed enrichment (X). Following normalization, enrichment scores for each bin were determined using a Poisson framework: The maximum expected signal (k) within 5 kb, 10 kb, and the entire genome were calculated using a probability density function, the p value tied to a specific bin enrichment calculated using P(X > k). P-values were adjusted globally using the Benjamini-Hochberg method. The data workflow and processed files are available at github.com/VanBortleLab/NuRDcomplex_metamap.

### Chromatin-IP and ChIP-seq analysis

Equal numbers of THP-1 monocytes, THP-1 derived macrophages, and ML-60218 treated monocytes were collected (∼10 million cells per ChIP experiment) and resuspended in growth media at 1 × 10^6 cells/mL and cross-linked with rotation at room temperature in 1% formaldehyde for 10 min. Cross-linking was quenched with the addition of 200 mM glycine and an additional 5 min of rotation at room temperature. Cross-linked cells were then spun down and resuspended in 1× RIPA lysis buffer, followed by chromatin shearing via sonication (3 cycles using a Branson sonicator: 30 s on, 60 s off; 20 additional cycles on a Bioruptor sonicator: 30 s on, 30 s off). Individual ChIP experiments were performed on pre-cleared chromatin using antibody-coupled ChIP grade Protein G magnetic beads (Cell Signaling Technology). 5 ug of antibody per ChIP was coupled to 60 uL of beads and rotated overnight with sheared chromatin at 4°C. Beads were then washed 5× in ChIP wash buffer (Santa Cruz), 1× in TE, and chromatin eluted in TE + 1% SDS. Cross-linking was then reversed by incubation at 65°C overnight, followed by digestion of RNA (30 min RNase (Cell Signaling Technology cat #7012), 1mg/exp incubation at 37°C) and digestion of protein (30 min, 30ug/1 exp proteinase K (Cell Signaling Technology cat #10012) incubation at 45°C). ChIP DNA was then purified on a minElute column (Qiagen), followed by DNA library preparation (NEBNext Ultra II DNA Library Prep Kit for Illumina) and size selection of 350-550 bp fragments via gel extraction (Qiagen). Samples are quantified using Bioanalyzer and sequenced with Nova X 25B paired-reads 300-cycle on the NovaSeq X Plus platform. Fastq files were generated and demultiplexed with the bcl-convert v4.1.7 Conversion Software (Illumina).

Fastq files were generated and demultiplexed with the bcl2fastq v2.20 Conversion Software (Illumina). Raw paired-end reads were trimmed with Trim_Galore/0.6.5, and read quality was assessed using FastQC. Trimmed reads were thereafter aligned to the human reference genome GRCh38_noalt_as using Bowtie2 version 2.3.5, and resulting SAM files were converted to BAM, sorted, and indexed using SAMtools version 1.9. Normalized signal tracks (20 bp bin size, smoothed over 60 bp) were generated with deepTools version 3.2.1 using the bamCoverage command with RPGC normalization. Read coverage over RNA central noncoding genes (gene body +/-150 wbp). Custom-defined genomic windows were computed using BEDTools version 2.26.0 with the coverage function. Peaks were identified using MACS2 version 2.2.5. Statistical scoring of NuRD occupancy was performed as previously described. Feature dominance analyses were computed and visualized via the dominatR package (github.com/VanBortleLab/dominatR).

### Small RNA-seq, FastGRO, and analyses

Fastq files were generated and demultiplexed with the bcl2fastq v2.20 Conversion Software (Illumina). Total steady-state small RNA was purified from equal numbers (∼2 million cells per small RNA-seq experiment) of THP-1 monocytes ± Romidepsin exposure using the mirVana™ PARIS™ RNA and Native Protein Purification Kit (Catalog# AM1556, Invitrogen) according to the manufacturer’s protocol for sRNA enrichment. The small RNA were treated with Antartic Phosphatase and PNK and libraries were constructed with the NEBNext Small RNA Sample Prep kit (NEB).

FastGRO nascent RNA-sequencing experiments were performed according to a previously described protocol (www.protocols.io/view/fastgro-yxmvmxe4ol3p/v1) with the following modifications: 10 million THP-1 nuclei were used per experiment; freshly isolated nuclei were pretreated with 25nM final concentration ML-60218 (Catalog# 557403-10MG, SigmaAldrich) or DMSO for 10 min and in-vitro run-on was carried out with 25nM ML-60218 or DMSO. Purified small RNA were processed with Antartic Phosphatase and PNK, and libraries constructed with the NEBNext Small RNA Sample Prep kit (NEB). The libraries were sequenced on one 10B lane for 101 cycles from one end of the fragments on a NovaSeq X Plus with V1.0 sequencing kits.

Raw single-end reads were trimmed with Trim_Galore to remove adapter sequences and low-quality bases, and read quality was assessed using FastQC. Trimmed reads were aligned to the custom 350 active gens index using Bowtie2 version 2.3.5.1, and resulting SAM files were converted, sorted, and indexed as BAM files using SAMtools version 1.9. Genome-normalized coverage tracks were generated using deepTools version 3.2.1. Reads counts over RNA central ncRNA genes (gene body +/-50 bp) were extracted with BEDTools version 2.28.0 (coverage function, -counts mode) and used for global background normalization. Differential analyses were performed using DEseq2.

### Contact Insulation calculations

THP-1 Hi-C data were obtained through SRA bioproject number PRJNA385337. For each active Pol III gene, contact reads over a 400 kb region (± 200 kb from the gene center) were extracted at 5 kb resolution using JuicerTools version 1.22.01^72^. For each window, all bin pairs were enumerated and categorized relative to the gene center as upstream, downstream, or cross-domain (contacts bridging across the gene). Self-diagonal bins were excluded. Cross-domain contact strengths were compared to distance-matched background bins grouped by 50 kb intervals to control for contact decay. For each window, 1000 permutations were performed by random sampling of global background bins, and a two-sided Mann–Whitney U test was used to assess significance (SciPy). Observed and expected medians from the permuted distributions were used to compute z-scores and permutation-based p-values. Median diagonal contact values are recorded as local contact intensity.

### Agglomerative clustering of Pol III-transcribed genes

Adjacent active Pol III genes are grouped into the same cluster if the distance between the transcription start site (TSS) of the downstream gene and the transcription end site (TES) of the upstream gene is ≤20 kb. A cluster terminates when no additional active Pol III gene is found within 20 kb, and a new cluster begins at the next downstream gene.

### Dynamic range index calculations

To minimize bias, the read counts extracted from small RNA seq (described above) are normalized as read per million, then quantile normalized, then log10 transformed prior to resampling. Resampling are introduced to minimize influence by extremely abundant tRNAs: Within each sample, a permutation-based sampling approach (n = 5000) was applied: in each iteration, 80% of active tRNA genes were randomly selected from the dataset, and the average expression of the top five genes within that subset (i.e., the five highest quantile-normalized log_10_ RPM values) was computed. The mean, standard deviation (SD), and interquartile range (25th–75th percentiles) were then derived from the 5000 simulated values to represent the empirical dynamic range of small RNA abundance in each condition. To assess the significance of observed differences in dynamic range between Romidepsin-treated and DMSO control samples, a permutation-based null distribution was generated. For each gene, the normalized RPMs were shaffled into 3 new control and 3 new treated groups, the dynamic range of these groups are calculated as described above. Then the p value is calculated as p=n{ expected abs[mean(treated)-mean (control)] >= observed abs[mean(treated)-mean (control)] x / n{perm}

## Resource Availability

Mass spectrometry proteomics data are deposited through ProteomeXchange Consortium via the PRIDE^73^ partner repository with the dataset identifier PXD061197 and can be accessed using reviewer token PiWIYkcMYF8A. All raw and processed ChIP-seq, ATAC-seq, small RNA-seq data are available through NCBI GEO accession GSE310006, and can be accessed using secure token uhmpekqsxzonzmp. FastGRO nascent RNA-seq data are available through NCBI GEO accession GSE311392, and can be accessed using secure token sxqrcygkflwdryj.

## Acknowledgements

We thank Drs. Alvaro Hernandez, Chris Wright, Danman Zhang, and staff at the Carver Biotechnology Center for sequencing services, and administrators of the Carl R. Woese Institute for Genomic Biology (UIUC) Biocluster for computational support. We thank Drs. Peter Yau, Justine Arrington, Brian Imai, and staff at the Carver Biotechnology Center for proteomics services. We thank members of the Van Bortle lab for helpful suggestions. This work was supported by the National Institutes of Health, National Human Genome Research Institute (NHGRI) grant R00HG010362 and Roy J. Carver Charitable Trust grant 26-6113 to KVB.

## Author contributions

Study design: RC, KVB. Data collection: RC, KVB, JY, SZ. Data analysis: RC, RKC, SL, KVB. Data interpretation: RC, KVB. Writing: RC and KVB with comments from all authors.

## Competing interest statement

The authors declare no competing interests

## Supplementary Figures

**Supplementary Figure 1.**
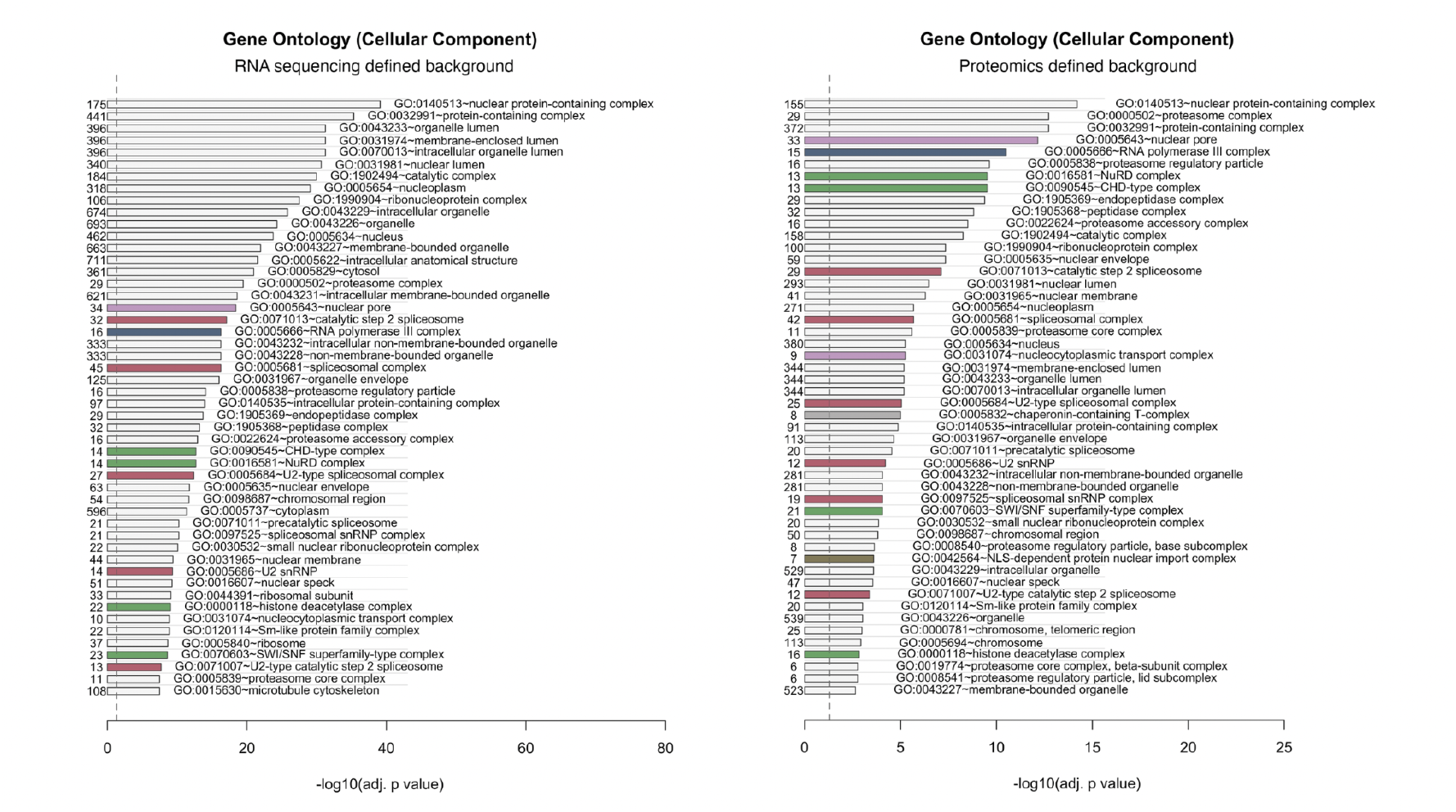
Gene ontology Cellular component enrichment analysis of Pol III interactome. Terms ranked by adjusted p-value. GO terms related to Figure 1a are highlighted with corresponding colors.

**Supplementary Figure 2.**
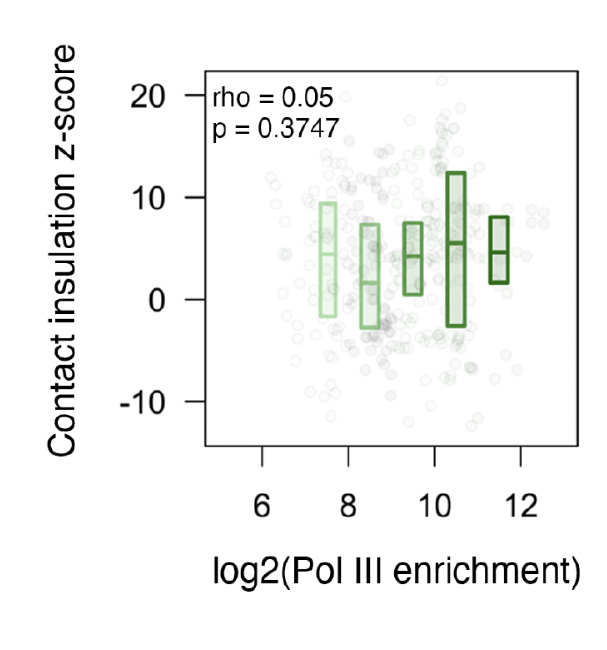
Pol III occupancy is not correlated with contact insulation. Distribution of contact insulation scores, measured by Hi-C, at increasing levels of Pol III enrichment.

**Supplementary Figure 3.**
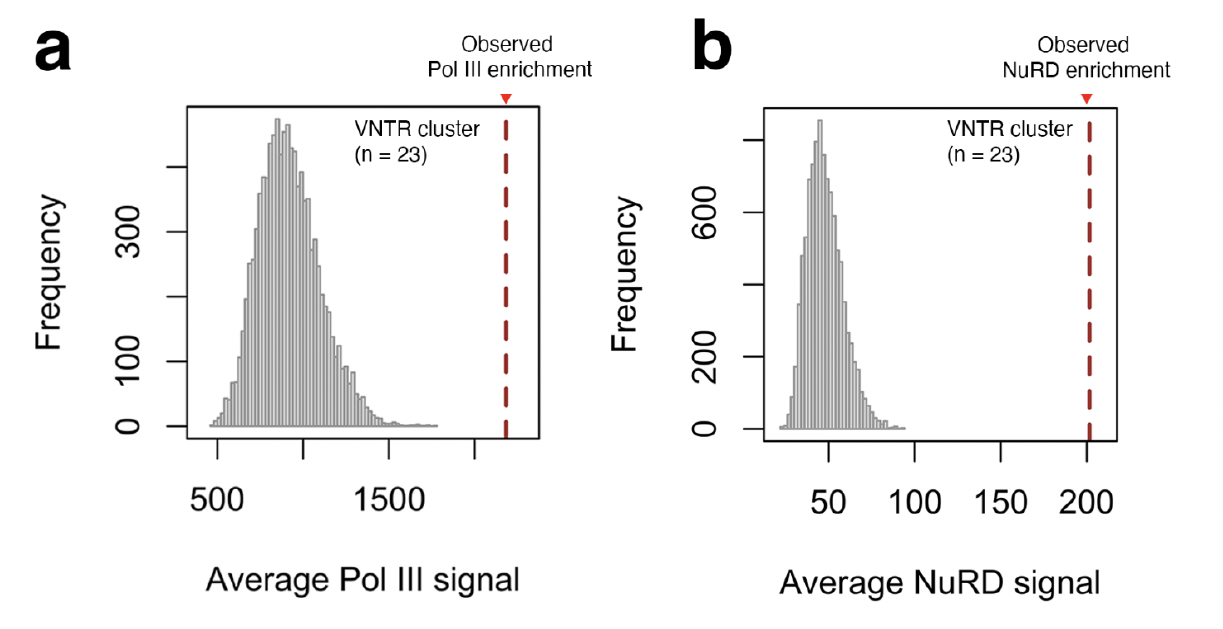
NuRD and Pol III enrichment peaks at VNTR genes. Histograms depict an empirical cluster-level null expectation.

**Supplementary Figure 4.**
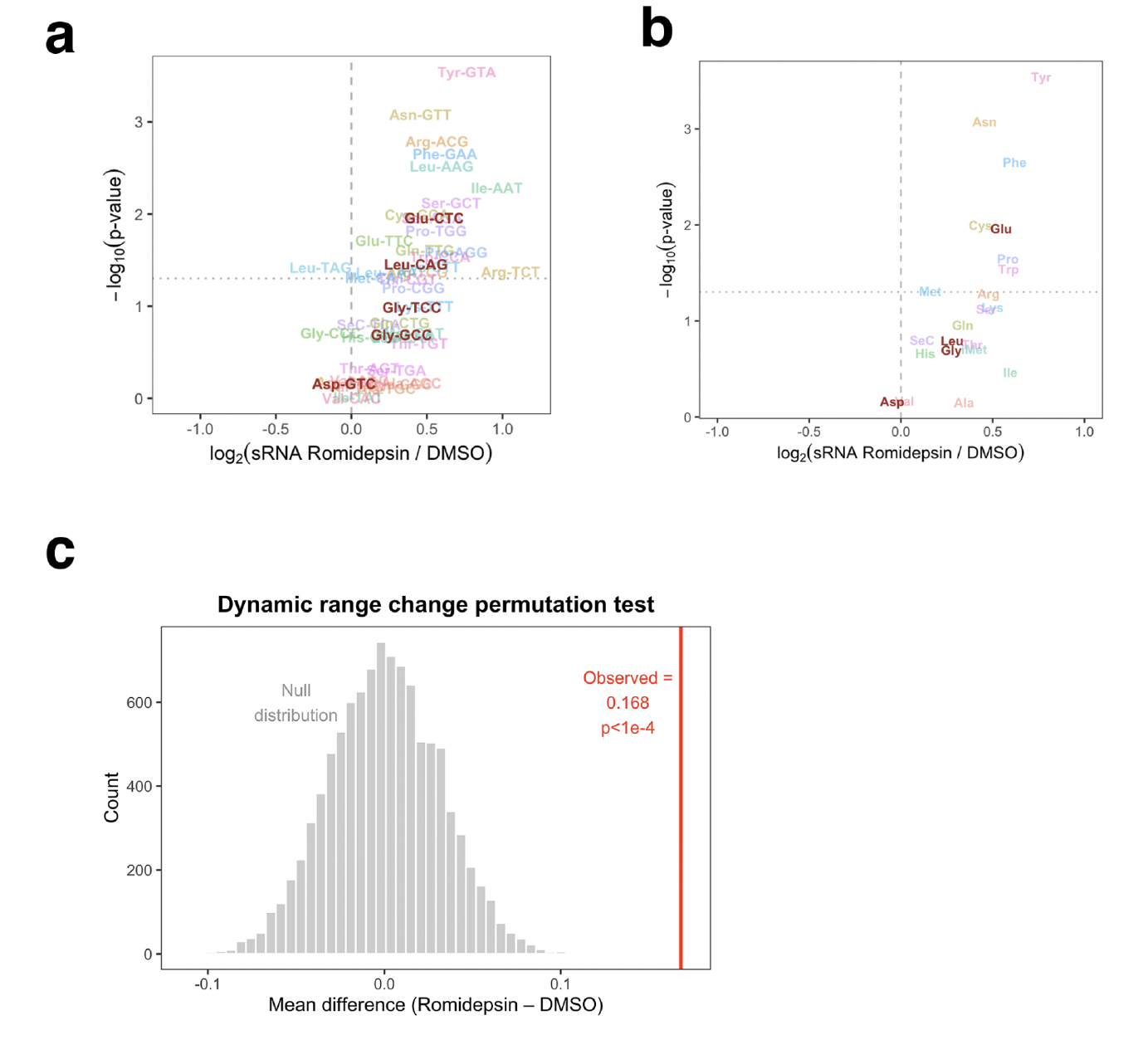
NuRD function antagonizes tRNA isoacceptor and isotype levels. **a)** Volcano plot showing differential small RNA-seq analysis at the tRNA isoacceptor level. **b)** Volcano plot showing differential small RNA-seq analysis at the tRNA isotype level. **c)** Histogram of permutation-based test results illustrating the change in Pol III dynamic range following Romidepsin treatment.

**Supplementary Figure 5.**
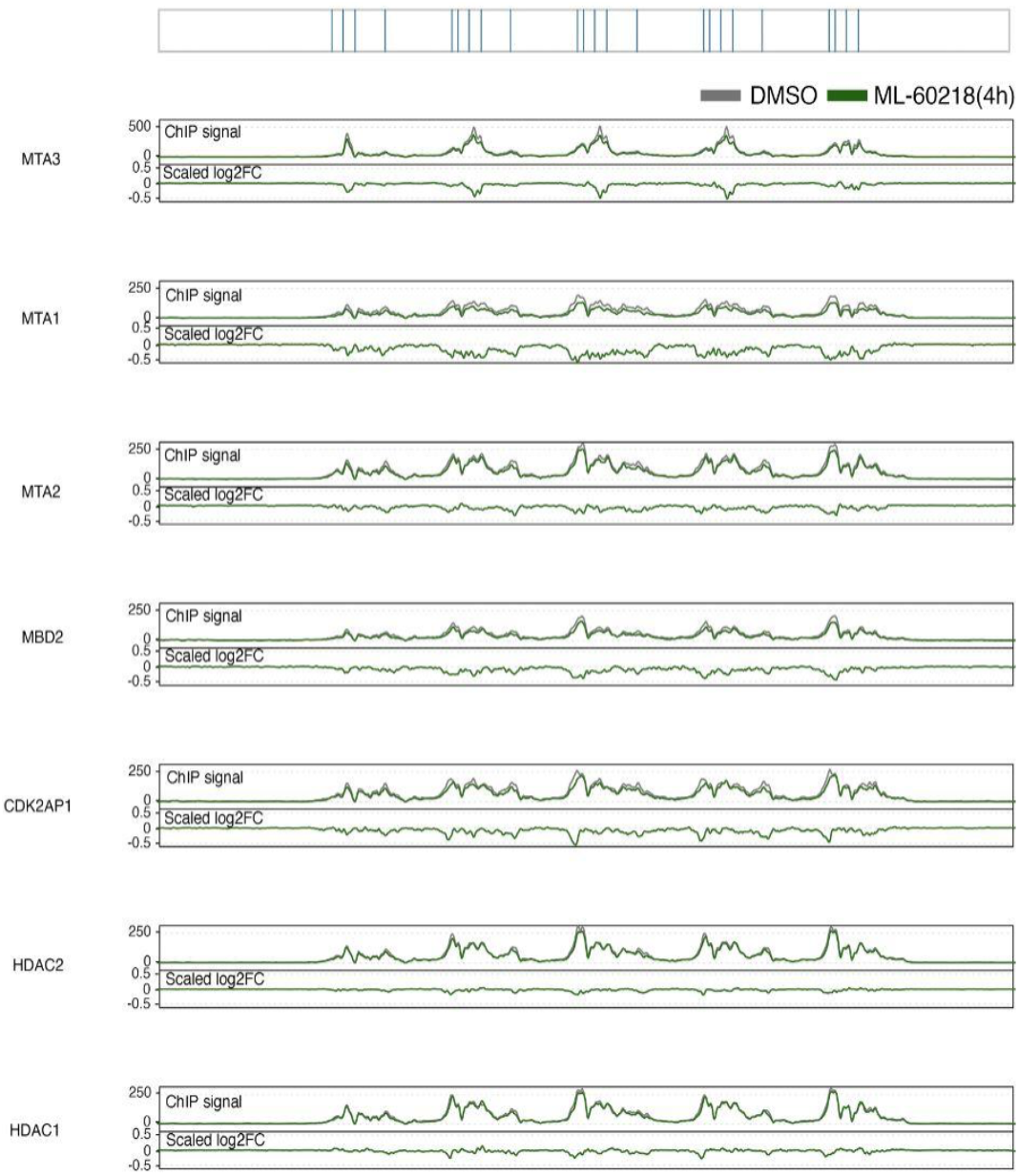
Pol III inhibition ChIP signal tracks and scaled log2 fold-change of NuRD subunits. **Top:** The blue vertical line marks the locus of the VNTR tRNA genes. **Bottom:** ChIP signal panels show RPGC-normalized enrichment in control (grey) and ML60218-induced Pol III inhibition (green). Scaled log2FC panels display log2(treated signal / control signal) × relative enrichment.

**Supplementary Figure 6.**
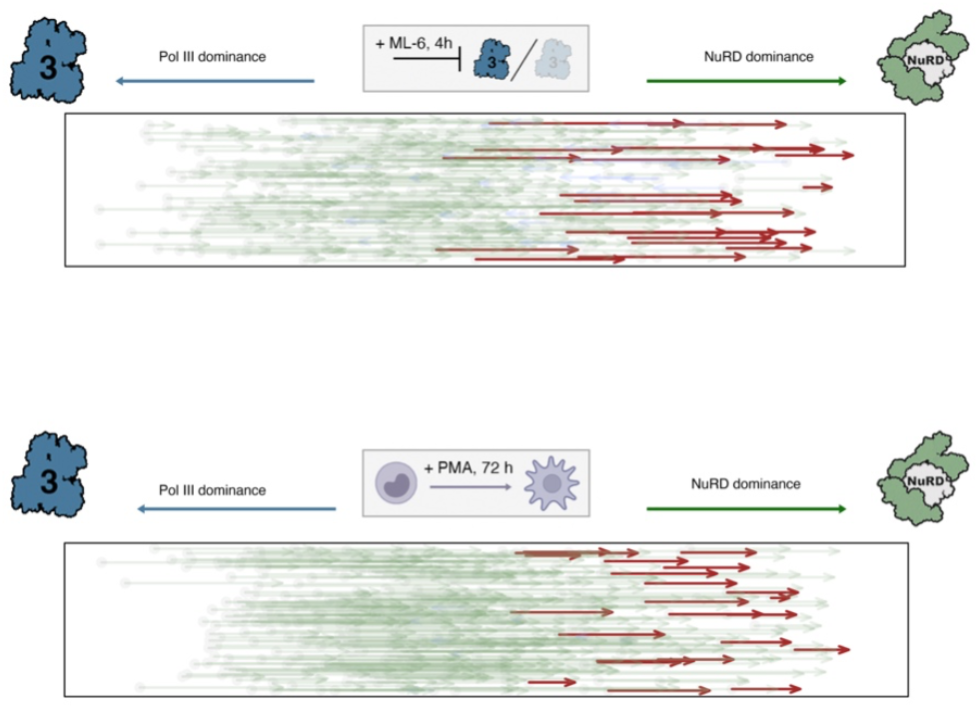
Gene-level dominance analysis comparing Pol III and NuRD-binding signatures in control and Pol III inhibition or differentiation. **Top**: Differential dominance analysis following ML6 induced Pol III inhibition relative Pol III occupancy defined by POLR3G occupancy. Genes with increased NuRD dominance colored in green, genes with increased Pol III dominance colored in blue, VNTR tRNA are highlighted in red. **Bottom:** Differential dominance analysis following PMA induced THP-1 monocyte differentiation, relative Pol III occupancy calculated by averaging POLR3B and POLR3D occupancy.

**Supplementary Figure 7.**
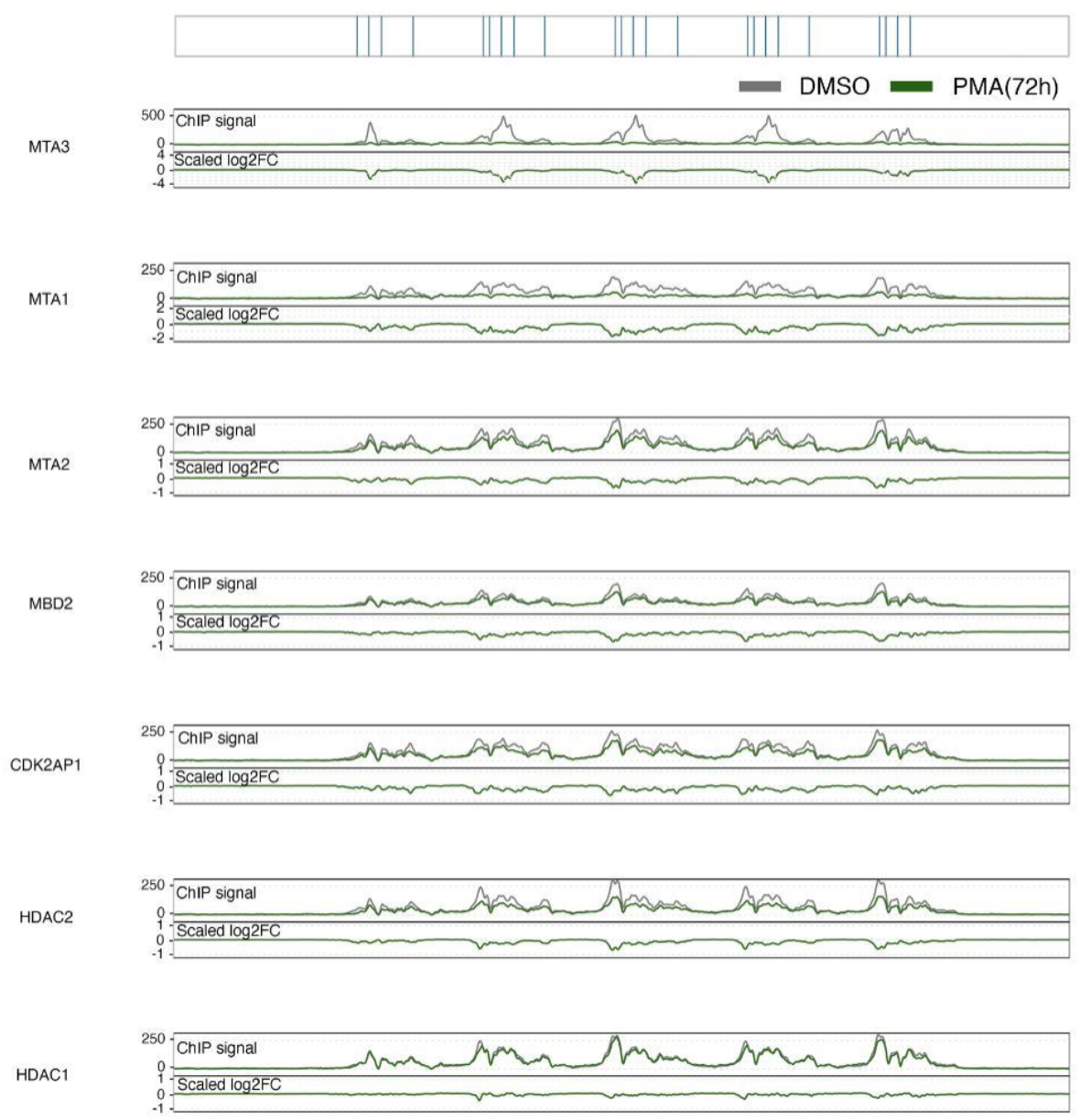
Monocyte differentiation ChIP signal tracks and scaled log2 fold-change of NuRD subunits. Top: The blue vertical line marks the locus of the VNTR tRNA genes. Bottom: ChIP signal panels show RPGC-normalized enrichment in control (grey) and PMA-induced differentiation (green). Scaled log2FC panels display log2(treated signal / control signal) × relative enrichment.

**Supplementary Table 1.**
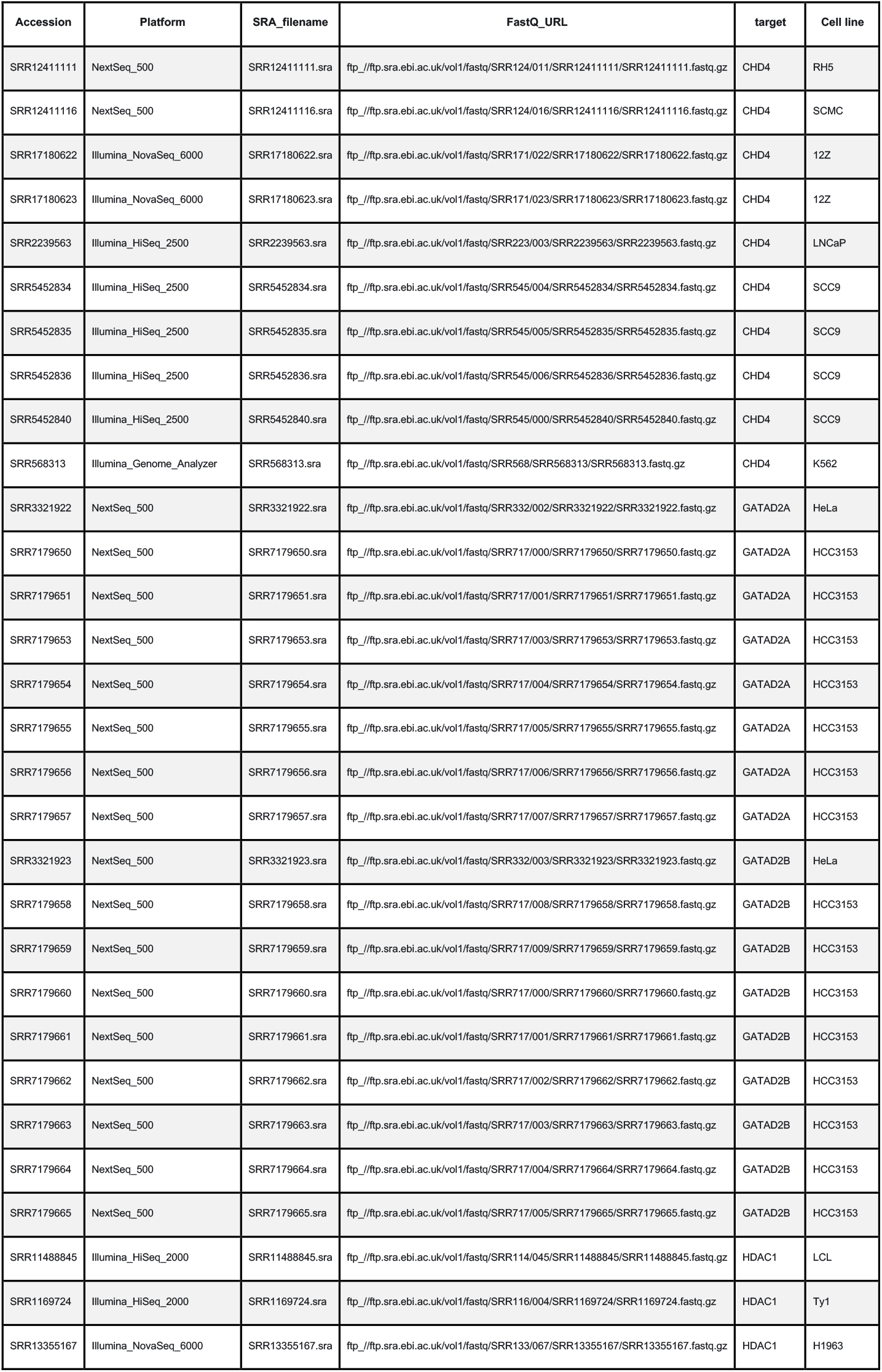
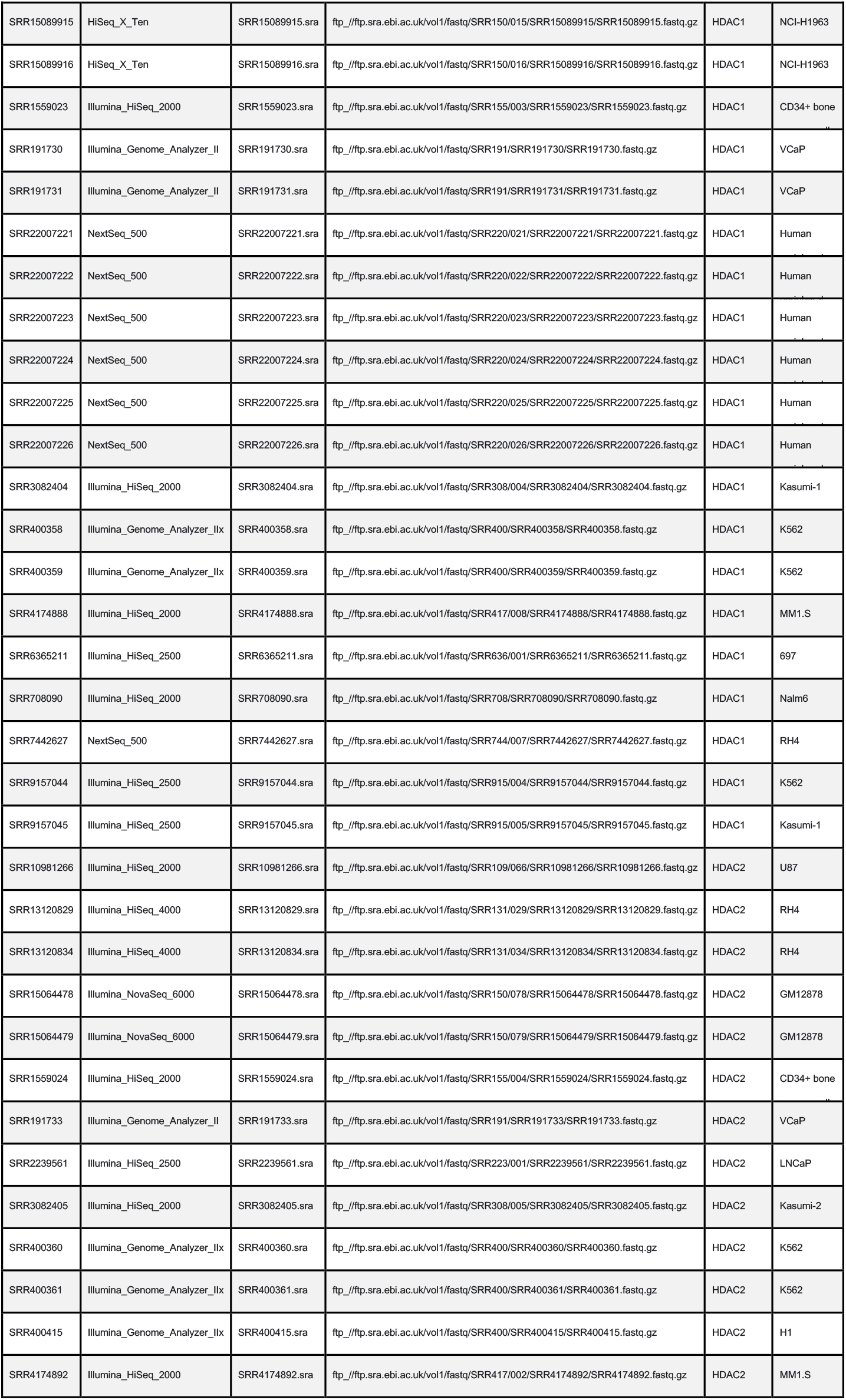
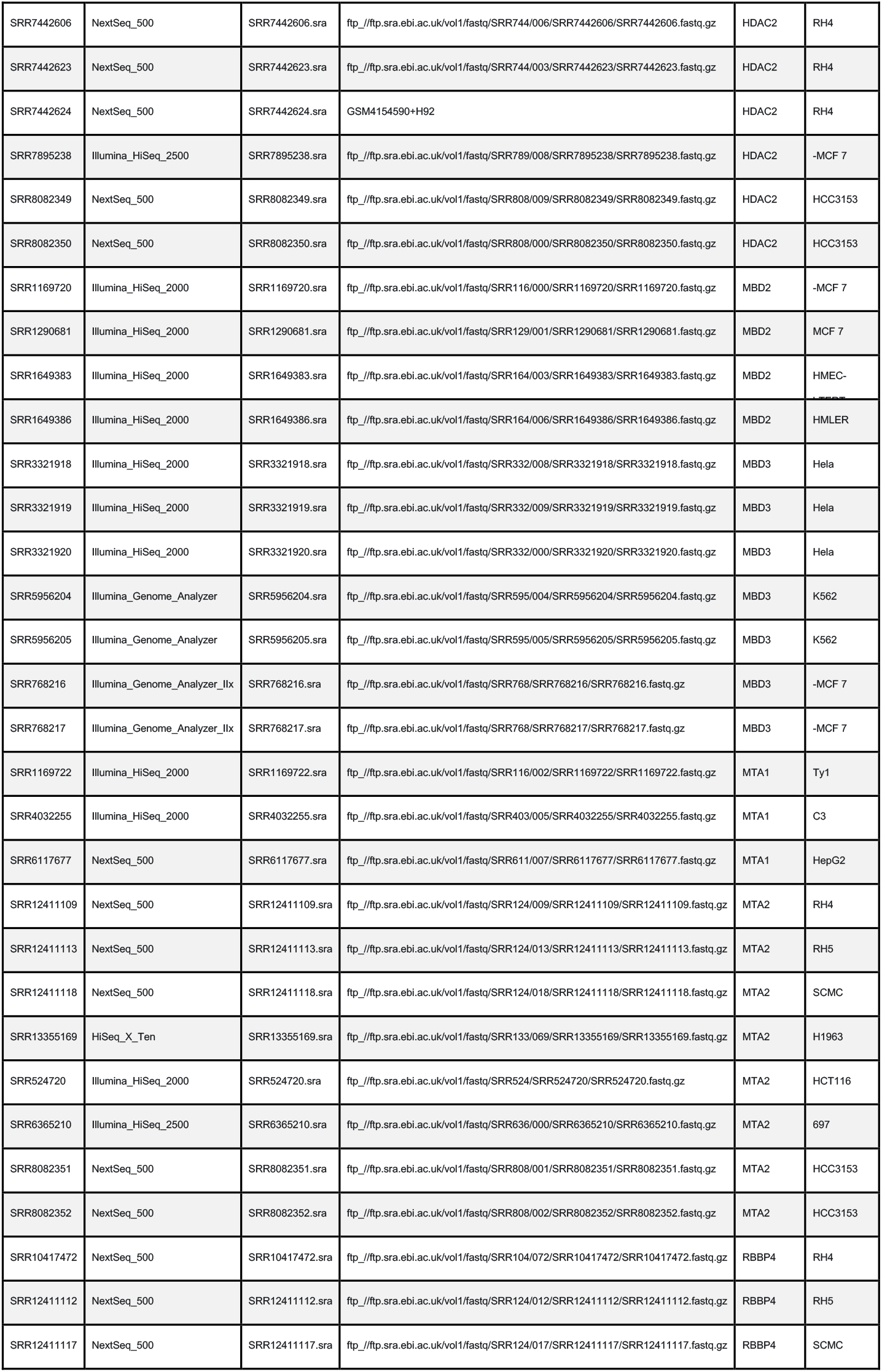

## Notes

### Competing Interest Statement

The authors have declared no competing interest.

